# Topology Matters: The Trade-off Between Wasserstein Critics and Discriminators for Single-Cell Data Integration

**DOI:** 10.1101/2025.11.12.688061

**Authors:** Kendall Reid, Genevieve Stein-O’Brien, Erhan Guven

**Author notes:** These authors contributed equally to this work (Genevieve Stein-O’Brien, Erhan Guven).

## Abstract

**Motivation:** Integrating single-cell RNA sequencing experiments (scRNA-seq) across technologies is hindered by severe technical batch effects that confound analysis and mask biological variation. Adversarial autoencoders are a popular solution to correct for these confounding effects, often relying on discriminator networks that approximate the Jensen-Shannon divergence. Previous research has established that the Jensen-Shannon divergence suffers from vanishing gradients when distributions do not overlap, a common phenomenon when datasets come from different sequencing technologies, leading to failed training. In contrast, the Wasserstein distance remains a valid metric with informative gradients even for disjoint distributions. While both approaches appear in the literature, no study has rigorously isolated the adversarial objective to systematically evaluate its impact on batch alignment, biological conservation, and scalability across varying dataset complexities.

**Results:** We introduce a multi-class reference-based Wasserstein critic to systematically benchmark adversarial objectives. We find that the Wasserstein critic yields superior mixing; however, extensive reference sensitivity analysis reveals that the Wasserstein critic is prone to over-correction resulting in collapsed cellular representations; that its integrative performance is dependent on a topologically dense reference batch; and that it scales poorly with the number of batches. In contrast, we find that the “weak” integration characteristic of discriminators acts as a protective measure against over-correction. By highlighting the trade-offs between these methods, we aim to empower researchers to choose the correct method for their specific needs.

**Availability and Implementation:** **Source code is available at** https://github.com/kreid415/wasserstein-critic-deconfounding. **Data are available at** https://figshare.com/articles/dataset/Benchmarking_atlas-level_data_integration_in_single-cell_genomics_-_integration_task_datasets_Immune_and_pancreas_/12420968/1.

## Introduction

High-throughput transcriptomic profiling provides high-resolution snapshots of cellular states, enabling the study of complex biological systems. However, these measurements are highly susceptible to systematic confounding that obscures true biological variation. These confounders arise primarily from technical batch effects—such as the sequencing technology utilized (Tung et al., 2017) or specific laboratory protocols—and are often complicated by underlying biological covariates, such as sample age (Stegle et al., 2015). This problem is compounded across experiments and laboratory conditions, making it impossible to compare gene expression results without controlling for these confounding variables (Leek et al., 2010; Goh et al., 2022; Yu et al., 2024).

Latent-space models trained with adversarial losses constitute a popular class of algorithms for removing confounding bias, operating by encouraging learned representations free of information from confounding variables (Dincer et al., 2020; He et al., 2025). These techniques typically rely on discriminator networks that predict the batch origin of a sample, thereby approximating the Jensen-Shannon (JS) divergence between batch distributions. A fundamental theoretical limitation of the JS divergence is that it fails to provide a meaningful gradient when two distributions have non-overlapping support (Arjovsky et al., 2017). In this scenario, a standard discriminator can perfectly separate the batches, leading to gradient saturation and training instability that prevents the generative model from successfully aligning the underlying data.

This instability has been addressed in the broader deep learning literature by replacing the discriminator with a Wasserstein critic—a network constrained to approximate the Earth Mover’s (Wasserstein) distance (Arjovsky et al., 2017). Unlike the JS divergence, the Wasserstein distance remains a continuous and differentiable metric even when distributions are non-overlapping, providing robust gradients that drive alignment regardless of the initial separation. While Wasserstein-based approaches have been proposed for biological batch correction (Wang et al., 2022, 2023), they often include additional steps that obscure the contribution of the adversarial mechanism. To date, a rigorous isolation of the adversarial mechanism—comparing the standard discriminator against a Wasserstein critic while holding the autoencoder backbone constant—has not been performed.

In this work, we strip away these auxiliary constraints to isolate the Wasserstein objective and investigate the specific contribution of the adversarial mechanism. To enable this rigorous comparison in realistic multi-batch settings, we implement a *multi-headed critic* formulation that generalizes the Wasserstein distance to multiple batches. This formulation allows us to explicitly and simultaneously minimize the Earth Mover’s distance between each source batch and a biologically grounded reference. By holding the autoencoder backbone constant, we control for architectural variables and directly test the efficacy of the objective function. We demonstrate that while replacing the standard Jensen-Shannon discriminator with the Wasserstein critic improves performance in disjoint settings, it introduces a topological bottleneck in scaling and a tendency to over-correct for batch effects, resulting in the loss of biological signal. Our results reveal a fundamental trade-off: the Wasserstein objective is a stronger integrator and is ideal for scenarios with severe, disjoint batch effects and overlapping cell types, whereas the discriminator scales more effectively in fragmented atlases with disjoint cell types or sensitive cellular boundaries.

## Related Work

The integration of experimental gene expression data remains a critical challenge in computational biology (Luecken et al., 2025), giving rise to a suite of solutions tailored to different analytical constraints (Tran et al., 2020). These methods can be broadly categorized into traditional statistical frameworks, geometric alignment heuristics, and deep generative models (Luecken et al., 2022). Traditional statistical methods, such as limma (Ritchie et al., 2015) and ComBat (Johnson et al., 2006), model expression data through linear frameworks to regress out confounding technical factors. Conversely, geometric methods rely on iterative alignment to map query datasets into a shared space, utilizing heuristics like Mutual Nearest Neighbors (MNN) (Haghverdi et al., 2018) or clustering-aware projections (Harmony (Korsunsky et al., 2019), Seurat (Stuart et al., 2019)).

More recently, deep generative models, such as the conditional variational autoencoder scVI (Lopez et al., 2018), have emerged to model the underlying non-linear data-generating process. Within this class, adversarial learning offers a powerful alternative, allowing for end-to-end differentiable deconfounding. Early adversarial approaches, such as AD-AE (Dincer et al., 2020), utilize standard discriminator networks to remove confounding information from the latent space. Recent works have begun to incorporate Wasserstein-based objectives, like ResPAN (Wang et al., 2022), which combines a Wasserstein critic with Random Walk Mutual Nearest Neighbors (RW-MNN) to guide alignment. Similarly, IMAAE (Wang et al., 2023) employs a Wasserstein-based adversarial autoencoder to tackle scenarios where cell types are disjoint across batches. Notably, Wang et al. explicitly attribute the failure of reference-based alignment in these settings to a “lack of constraints” when the anchor batch is sparse, motivating their decision to enforce a global Gaussian prior. Consequently, the specific contribution of the *Wasserstein critic itself* —as a stable replacement for the discriminator in pure adversarial deconfounding—has not been isolated or rigorously evaluated. In this work, we bridge this gap by stripping away these auxiliary additions and systematically comparing the Wasserstein critic against the standard discriminator within a controlled, state-of-the-art autoencoder framework.

## Materials and Methods

To rigorously isolate the impact of the adversarial objective, we employ a fixed Variational Autoencoder (VAE) backbone from scCRAFT (He et al., 2025). We hold the encoder, decoder, and auxiliary loss functions constant across all experiments, varying only the adversarial regularization mechanism (standard discriminator vs. Wasserstein critic). The total objective consists of four components: (1) a reconstruction loss, (2) a prior regularization loss, (3) a triplet loss to preserve local structure, and (4) the adversarial deconfounding loss. A simplified architecture is shown in Figure 1.

**Figure 1.**
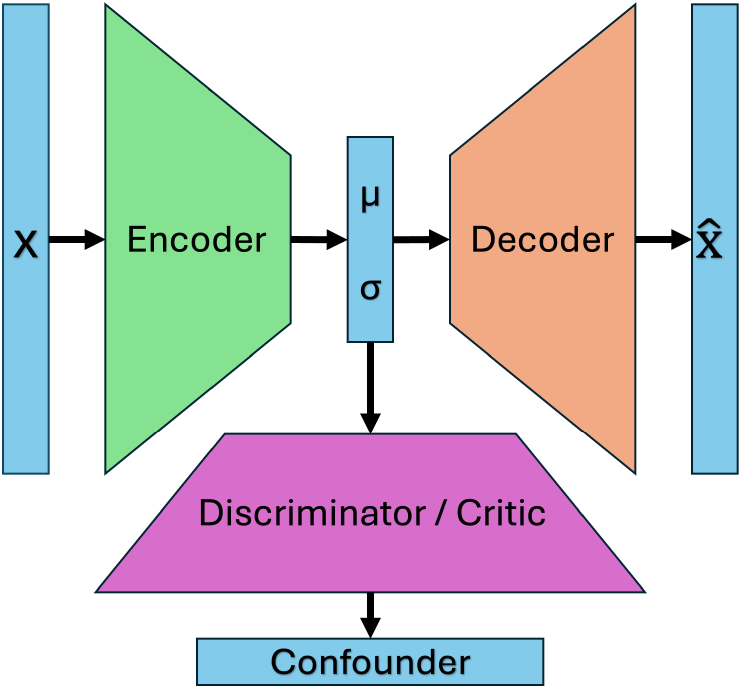
Experimental Architecture. To isolate the impact of the adversarial objective, we employ a fixed VAE backbone. We systematically swap the adversarial regularizer between a standard discriminator and Wasserstein critic to compare their efficacy in removing confounding information.

### Variational Autoencoder and Reconstruction Loss

The model takes log1p-transformed gene expression data, 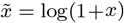, as input and aims to reconstruct the original count values, *x*. To account for the sparsity of genomic data, the model assumes the target data *x* follows a negative binomial (NB) distribution. The encoder network, 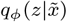, is a two-layer feedforward network that maps the input vector 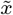 to a vector of size 512, 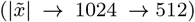. This vector is fed into parallel linear layers that output the mean (*µ*) and variance (*σ*^2^), each of size 256, which parameterize the latent normal distribution *Z* ~ *N*(*µ, σ*^2^). Within the encoder, batch normalization layers and ReLU activations are applied after the hidden dense layers to ensure training stability.

The decoder network reconstructs the count-based expression profile by combining the sampled latent code *z* ~ *Z* with a one-hot encoded vector representing the known batch variable ID, *v*. This is achieved via two parallel networks *G*_1_ and *G*_2_:

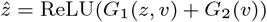

where *G*_1_ maps the concatenated latent code and batch variable code back to gene space (*z*||*v* → 512 → 1024 → |*x*| where || is the concatenation operator), and *G*_2_ captures baseline batch-specific effects (*v* → 512 → 1024 → |*x*|). This combined representation 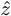 is fed into a final linear layer, followed by an exponential activation function, to predict strictly positive NB parameters for each gene (mean *µ* and dispersion *r*).

The reconstruction objective combines the NB negative log-likelihood with a cosine similarity loss to prevent the model from overfitting to high-count genes. To ensure stable optimization gradients across varying dataset sizes, the negative log-likelihood is computed as the mean across all genes *G*, rather than the sum:

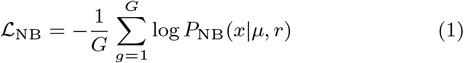

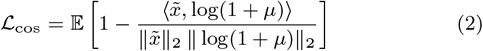

where ⟨·, ·⟩ denotes the dot product between the log-transformed original and the log-transformed reconstructed gene expression profiles.

To mitigate latent over-correction, the backbone model includes a dual-resolution triplet loss. To ensure training remains unsupervised and to prevent data leakage, we approximate cell types using pre-computed, two-tier Leiden clustering (coarse and fine resolutions, set empirically to 1.0 and 7.0, respectively) derived from the unintegrated data. For a given cell’s latent embedding acting as an anchor (*z*_*a*_), we initially sample positive (*z*_*p*_) and negative (*z*_*n*_) pairs based on the coarse cluster assignments. Crucially, to enforce strict local manifold preservation, we apply a high-resolution filter: if the anchor and the positive sample do not also share the same fine-resolution cluster assignment, the pair is discarded from the loss calculation. This objective enforces that cells of the strict same type remain closer in latent space than cells of different types:

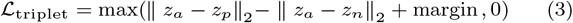

where the margin is set empirically to 5 as per the original implementation.

Equations 1, 2, and 3 are summed with the Kullback-Leibler (KL) divergence, which regularizes the latent space toward a standard normal prior to give the VAE-backbone:

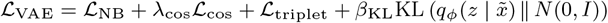

In our experiments, we set the hyperparameters *λ*_cos_ = 20 and *β*_KL_ = 0.005, identical to the original implementation (He et al., 2025).

### Baseline: Discriminator-Based Deconfounding

To serve as a baseline for comparison, we implement a standard adversarial deconfounding approach using a discriminator network *h*_*υ*_ : Ƶ → 𝒱, that attempts to predict the confounding variable *v* ∈ 𝒱 (e.g., batch ID) from the latent embedding code *z* ∈ Ƶ. The discriminator is trained to minimize the prediction error (typically cross-entropy), effectively approximating the Jensen-Shannon divergence between confounding distributions from the data source, 𝒟:

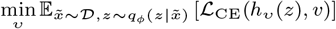

where ℒ_CE_ is the cross-entropy loss.

The discriminator architecture consists of a three-layer Multi-Layer Perceptron (MLP) mapping from the sampled latent codes to a vector of probabilities for each batch (|*z*| = 256 → 128 → 128 → |𝒱|), with ReLU activations following the first two layers. The final linear layer outputs raw logits, which are subsequently passed through a log-softmax operation within the cross-entropy objective.

### Multi-Headed Wasserstein Critic Formulation

To avoid the quadratic computational cost (*O*(|𝒱|^2^)) of pairwise matching in multi-batch settings, we formulate a *reference-based multi-headed critic*. We designate one batch as the *reference* (*r*) and align all other source batches (*i* ≠ *r*) to this biological ground truth. To measure the impact of reference choice, we perform sensitivity analysis by training with each batch as reference and compare the variance in results.

We modify the critic network *C*_*ρ*_ : Ƶ → ℝ^|𝒱|^ to output a vector of scores. The *i*-th component, *C*_*ρ*_(*z*)_*i*_, outputs the Kantorovich potential (critic score) required to estimate the Wasserstein distance between batch *i* and the reference batch. The critic is trained to maximize the estimated Wasserstein distance between the reference and source batches, finding the tightest Kantorovich-Rubinstein bound. In practice, this is implemented by minimizing the negative of the distance, regularized by a gradient penalty across all non-reference batches:

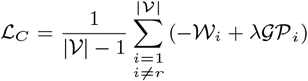

where 𝒲_*i*_ represents the unconstrained Wasserstein distance estimation and 𝒢𝒫_*i*_ is the gradient penalty for batch *i*:

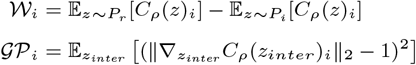

where *λ* is the gradient penalty coefficient (set to 10 in our experiments). To enforce the 1-Lipschitz constraint required by the Wasserstein-1 metric, we apply a Gradient Penalty (GP) (Gulrajani et al., 2017) individually for each head. This penalty is evaluated on samples *z*_*inter*_ interpolated uniformly along straight lines between the latent representations of the reference batch and the specific non-reference batch *i*: *z*_*inter*_ = *ϵz*_*i*_ + (1 − *ϵ*)*z*_*r*_, with *ϵ* ~ *U* [0, 1]. We calculate this penalty on points interpolated strictly between the source batch and the reference batch to enforce the constraint along the relevant transport manifold. Crucially, to ensure this penalty remains well-defined, we employ a stratified sampling strategy that forces every mini-batch to contain samples from the reference distribution. This prevents the instability that arises if a mini-batch randomly lacks reference samples.

Simultaneously, the encoder network acts as the generator in this minimax game. While the critic maximizes the distance, the generator seeks to minimize it, thereby aligning the non-reference batches with the reference manifold. Because the generator has no influence over the reference batch distribution *P*_*r*_ or the gradient penalty, it minimizes the Wasserstein distance strictly by maximizing the critic’s score for its encoded source samples (*P*_*i*_). Consequently, the generator’s adversarial objective simplifies to:

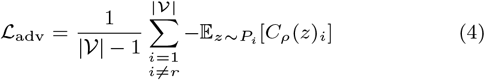

By minimizing this term, the encoder forces the latent representations of the disjoint source batches to become indistinguishable from the reference manifold under the evaluation of the critic.

We use a 10:1 (critic:encoder) update ratio, as the Wasserstein objective provides stable gradients even near optimality.

### Final Objective

To evaluate the impact of the Wasserstein objective, we define two distinct total loss functions for the encoder network. In both configurations, the encoder optimizes the fixed backbone losses (ℒ_VAE_), but the adversarial penalty varies depending on the chosen framework.

For the baseline discriminator model, the encoder acts as a standard generator, minimizing the negative cross-entropy to maximize classification error and successfully confuse the discriminator:

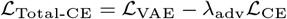

Conversely, for the Wasserstein critic formulation, the encoder acts as the generator, minimizing its specific adversarial component (Eq 4) to conceptually minimize the Wasserstein distance and align the latent distributions. Thus, the total encoder objective becomes strictly additive:

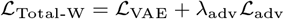

### Implementation, Training, and Model Selection

To evaluate performance across varying degrees of topological complexity and batch overlap, we utilized three standard benchmark datasets (Luecken et al., 2022): **Pancreas** (9 batches), **Lung** (2 batches), and **Immune** (4 batches). All models were implemented in PyTorch and trained on a single NVIDIA RTX 4090 GPU using the Adam optimizer (learning rate = 0.001, *β*_1_ = 0.5, *β*_2_ = 0.9); the *β*_1_ momentum parameter was intentionally reduced to stabilize adversarial dynamics. Models were trained with a batch size of 1024. Due to the multiple local updates required to enforce the Lipschitz constraint (10 critic updates per generator step), the Wasserstein-based models required approximately 24% more training time per epoch than the baseline discriminator (see detailed breakdown in Supplementary Figure S1 and Supplementary Table S1).

To ensure stability, the Gradient Penalty weight was fixed at *λ*_*GP*_ = 10 (Gulrajani et al., 2017), and we employed a warmup period for the first 10 epochs where *λ*_adv_ = 0 and the adversary’s weights were frozen. This allows for valid autoencoder reconstruction before adversarial alignment. We utilized class-balanced sampling with replacement across all experiments; this ensured equal representation of all source batches within each mini-batch, which both prevented mode collapse in smaller experimental batches and guaranteed well-defined interpolations for the Gradient Penalty. To prevent combinatorial explosion during training, triplet mining was capped at a maximum of 15 random positive pairs per cell-type per batch within each training step. Models were trained and evaluated using a nested cross-validation scheme (5 outer folds, 3 inner folds), where inner loops ran for 40 epochs and outer loops for 80 epochs. Hyperparameters were systematically selected to maximize the composite integration score, (iLISI − 10 * cLISI), on held-out data, where the cLISI penalty is scaled by a factor of 10 to balance the magnitude of the respective metrics.

## Experiments and Datasets

We conducted three targeted experiments to assess performance across varying levels of integration complexity:

1. **Binary Integration (Fundamental Baseline):** We first restricted the problem to the two largest batches in each dataset, simplifying the multi-class problem to a foundational binary classification task.
2. **Multi-Batch Scalability:** To understand how the adversarial objectives scale with problem complexity, we incrementally increased the number of source batches from 2 to |𝒱| (adding batches in descending order of size). The reference batch for these experiments was fixed to the largest batch in each respective dataset.
3. **Reference Sensitivity Analysis:** To evaluate the robustness of the multi-headed critic formulation, we exhaustively trained models using every possible batch as the reference target. This experiment determines whether the critic’s integrative performance is strictly dependent on the specific topology and cellular density of the chosen reference.

### Confounding Scenarios

For every experiment, we evaluated two distinct confounding scenarios to test robustness against covariate correlation:

- **Balanced (Ideal):** We restricted the datasets to the intersection of shared cell types across all batches. In this scenario, batch and cell-type distributions are independent, representing an idealized integration task without biological confounding.
- **Unbalanced (Realistic):** We utilized the raw datasets where cell type proportions vary significantly between batches. This introduces a strong correlation between batch and underlying biology, a scenario known to drive severe over-correction and the ablation of biological signal in standard adversarial deconfounding.

For every combination of dataset, experiment, and scenario, we trained both a discriminator-based and a Wasserstein critic-based VAE and evaluated performance on held-out test data. For statistical comparisons, we applied the corrected paired *t*-test (Nadeau and Bengio, 2003) to account for the variance introduced by cross-validation overlap.

### Data Preprocessing

Prior to downstream analysis, we applied standard quality control filters, removing cells expressing fewer than 300 genes and genes expressed in fewer than 5 cells. Feature selection was performed by identifying the top 2,000 highly variable genes (HVGs) using the standard Seurat dispersion method. To satisfy the dual-input requirements of our architecture, the raw count matrices for these selected HVGs were retained as the reconstruction target (*x*) for the negative binomial loss. For the encoder input 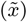, these counts were library-size normalized (scaled to 10^4^ reads per cell) and log1p-transformed (log(*x* + 1)).

A summary of the dataset compositions, including the total number of cells and distinct sequencing technologies (batches), is provided in Table 1. A comprehensive breakdown is available in Supplementary Table S2.

**Table 1.** Characteristics of the three scRNA-seq benchmark datasets. The table details the batch composition for the Pancreas, Lung, and Immune datasets, including the specific sequencing technology or protocol, total cell count, and distinct cell type diversity per batch. These datasets span a wide range of protocols, establishing a highly heterogeneous testbed for evaluating batch effect correction.

| Pancreas Dataset (9 Batches) |  |  | Lung Dataset (2 Batches) |  |  | Immune Dataset (4 Batches) |  |  |
| --- | --- | --- | --- | --- | --- | --- | --- | --- |
| Tech | Cells | Types | Protocol | Cells | Types | Chemistry | Cells | Types |
| CEL-Seq | 1,004 | 13 | 10x Chromium | 22,771 | 16 | 10x | 8,829 | 10 |
| CEL-Seq2 | 2,285 | 13 | Drop-Seq | 6,485 | 11 | Smart-seq2 | 1,022 | 4 |
| Fluidigm C1 | 638 | 13 |  |  |  | 10x Chromium v2 | 12,928 | 16 |
| inDrop-Seq V1 | 1,937 | 14 |  |  |  | 10x Chromium v3 | 10,727 | 12 |
| inDrop-Seq V2 | 1,724 | 14 |  |  |  |  |  |  |
| inDrop-Seq V3 | 3,605 | 14 |  |  |  |  |  |  |
| inDrop-Seq V4 | 1,303 | 14 |  |  |  |  |  |  |
| SMARTer | 1,492 | 4 |  |  |  |  |  |  |
| Smart-Seq V2 | 2,394 | 13 |  |  |  |  |  |  |

### Evaluation Framework

To strictly quantify the trade-off between removing confounding artifacts and preserving biological signal, we utilize a suite of complementary metrics. We categorize these into local and global measures to ensure our evaluation captures both fine-grained manifold alignment and macro-scale cluster structure. Before metric calculation, the 256-dimensional latent embeddings were reduced to 50 principal components using PCA to remove residual noise and standardize the dimensionality for distance calculations.

#### Local Metrics (Primary)

We prioritize the Local Inverse Simpson’s Index (LISI) (Korsunsky et al., 2019) as our primary evaluation tool. Unlike global summary statistics, LISI provides a cell-specific measure of integration quality, allowing us to assess how well the model aligns distributions within the local neighborhood of each cell:

- **iLISI (Integration):** Measures the effective number of batches in a cell’s local neighborhood. Like cLISI, we scale this metric to the interval [0, 1] via the transformation 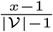, where |𝒱| is the total number of batches. Higher values indicate superior mixing of distinct experimental batches.
- **cLISI (Conservation):** Measures the effective number of cell types in a neighborhood. We scale this metric to the interval [0, 1] via the transformation 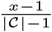, where *x* is the raw LISI score and |𝒞| is the total number of unique cell types in the dataset. Consequently, lower values indicate superior conservation, signifying that distinct cell types remain separated and are not erroneously merged.

LISI metrics were calculated with a perplexity of 30 and *k* = 90 nearest neighbors (Korsunsky et al., 2019).

#### Global Metrics (Secondary)

To assess macro-scale alignment, we employ standard metrics calculated using the scib Python framework (Luecken et al., 2022). To ensure consistent interpretability, we utilize the scaled versions of these metrics, where *higher values always indicate better performance*:

- **ASW Batch (Integration):** The Batch Silhouette Width measures the degree of batch mixing. We report the absolute scaled value, 1 − |ASW_batch_|, normalized to the interval [0, 1]. A score of 1 indicates perfect batch mixing, while 0 indicates complete separation.
- **ASW Cell Type (Conservation):** The Biological Silhouette Width measures the compactness of cell-type clusters. We report the standard silhouette width scaled to [0, 1], where 1 indicates perfectly distinct cell types and 0 indicates overlapping biological populations.
- **ARI (Adjusted Rand Index):** Measures the agreement between latent clustering and true cell-type labels (0 indicates random assignment; 1 indicates a perfect match).
- **PAGA Topology Preservation (Conservation):** Quantifies the preservation of the original biological manifold. We establish a baseline biological topology by computing the Partition-based Graph Abstraction (PAGA) (Wolf et al., 2019) connectivity matrix exclusively on unconfounded, single-technology batches in the raw PCA space. We then calculate the correlation between this baseline matrix and the PAGA matrix derived from the integrated latent space. Because neural networks non-linearly compress latent dimensions, absolute linear distances are inherently warped; therefore, we utilize the Spearman rank correlation coefficient (*r*) to accurately assess the preservation of the monotonic rank-order of biological neighborhoods. A higher score (→ 1) indicates the model successfully shielded the natural biological topology from artificial geometric collapse.

#### Generalization to Unseen Data

A critical component of our study is evaluating whether the adversarial alignment generalizes to new data. Unlike standard benchmarks that often evaluate on the training set, we compute all reported metrics on *held-out test data*. Specifically, we project test cells into the learned latent space and construct a joint *k*-nearest neighbor graph containing both training and test cells. Metrics are then computed specifically for the test nodes, measuring how well unseen query cells integrate into the established reference manifold.

## Results

### The Wasserstein Critic Achieves Superior Local Integration at the Cost of Biological Conservation

We first explore the utility of the Wasserstein critic in the simplest deconfounding scenario: integrating two distinct batches. This reduces the problem to binary classification and the corresponding binary Wasserstein formulation. We restricted the analysis to the two largest batches for each dataset (Pancreas, Immune, and Lung) under both balanced and unbalanced confounding scenarios. Hyperparameters were optimized using nested cross-validation, and performance was evaluated using six metrics covering both integration and biological conservation (Figure 2). Statistical significance was assessed using corrected paired t-tests (Nadeau and Bengio, 2003) (Table 2).

**Table 2.** Statistical comparison of the Wasserstein Critic and baseline Discriminator across balanced and unbalanced integration scenarios. The table reports the mean and standard deviation (SD) of various integration (iLISI, ASW Batch) and biological preservation metrics (cLISI, ASW Cell, ARI, PAGA Spearman) for the Pancreas, Immune, and Lung datasets. P-values represent the significance of the difference between the two models, corrected for cross-validation overlap (Nadeau & Bengio, 2003).

| Dataset | Metric | Balanced Scenario |  |  | Unbalanced Scenario |  |  |
| --- | --- | --- | --- | --- | --- | --- | --- |
|  |  | Critic (SD) | Discrim (SD) | p-value | Critic (SD) | Discrim (SD) | p-value |
| Pancreas | iLISI | 0.736 (0.022) | 0.620 (0.036) | 0.0064 | 0.740 (0.038) | 0.648 (0.029) | 0.0987 |
|  | ASW Batch | 0.894 (0.022) | 0.930 (0.017) | 0.0014 | 0.895 (0.014) | 0.932 (0.017) | 0.0141 |
|  | cLISI | 0.027 (0.007) | 0.010 (0.005) | 0.0138 | 0.024 (0.008) | 0.012 (0.005) | 0.1739 |
|  | ASW Cell | 0.570 (0.009) | 0.604 (0.006) | 0.0224 | 0.570 (0.009) | 0.605 (0.010) | 0.0012 |
|  | ARI | 0.670 (0.057) | 0.942 (0.018) | 0.0017 | 0.692 (0.113) | 0.938 (0.019) | 0.0399 |
|  | PAGA Spearman | 0.417 (0.090) | 0.506 (0.098) | 0.4402 | 0.410 (0.086) | 0.545 (0.073) | 0.0570 |
| Immune | iLISI | 0.713 (0.012) | 0.603 (0.013) | 4.84e-04 | 0.689 (0.030) | 0.562 (0.008) | 0.0040 |
|  | ASW Batch | 0.853 (0.021) | 0.872 (0.013) | 0.2607 | 0.873 (0.013) | 0.878 (0.008) | 0.5216 |
|  | cLISI | 0.035 (0.004) | 0.019 (0.003) | 0.0113 | 0.030 (0.002) | 0.016 (0.003) | 0.0012 |
|  | ASW Cell | 0.553 (0.003) | 0.571 (0.004) | 0.0073 | 0.517 (0.008) | 0.566 (0.003) | 0.0024 |
|  | ARI | 0.447 (0.030) | 0.556 (0.006) | 0.0051 | 0.439 (0.028) | 0.554 (0.007) | 0.0062 |
|  | PAGA Spearman | 0.459 (0.070) | 0.615 (0.083) | 0.1479 | 0.524 (0.071) | 0.559 (0.049) | 0.3811 |
| Lung | iLISI | 0.622 (0.041) | 0.498 (0.021) | 0.0175 | 0.455 (0.016) | 0.370 (0.024) | 0.0080 |
|  | ASW Batch | 0.870 (0.006) | 0.910 (0.007) | 0.0026 | 0.861 (0.011) | 0.898 (0.009) | 0.0212 |
|  | cLISI | 0.034 (0.004) | 0.017 (0.003) | 0.0031 | 0.025 (0.003) | 0.017 (0.001) | 0.0253 |
|  | ASW Cell | 0.551 (0.004) | 0.578 (0.005) | 8.63e-04 | 0.549 (0.006) | 0.566 (0.005) | 0.0290 |
|  | ARI | 0.473 (0.021) | 0.534 (0.009) | 0.0315 | 0.607 (0.074) | 0.565 (0.059) | 0.5359 |
|  | PAGA Spearman | 0.298 (0.031) | 0.526 (0.035) | 1.22e-04 | 0.554 (0.064) | 0.657 (0.055) | 0.2536 |

**Figure 2.**
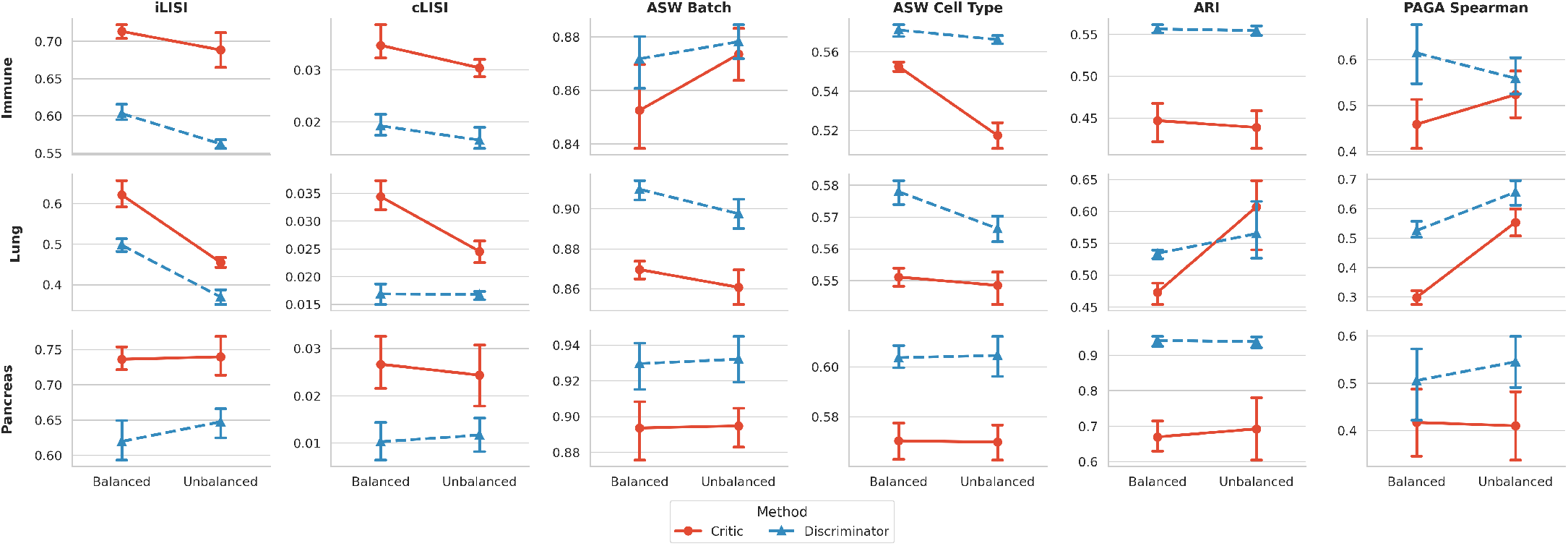
Performance comparison of adversarial deconfounding methods across datasets and batch scenarios. We evaluated the Wasserstein critic (Red circles, solid line) against a standard discriminator (Blue triangles, dashed line) across three datasets (Pancreas, Immune, Lung). Performance was assessed using six metrics. **Integration metrics:** iLISI and ASW Batch (higher is better) measure the removal of batch effects. **Conservation metrics:** ASW Cell Type, ARI, and PAGA Spearman (higher is better) and cLISI (lower is better) measure the retention of biological signal. The x-axis denotes the confounding scenario (Balanced vs. Unbalanced). Points represent the mean score across 5-fold cross-validation, and error bars indicate the 95% confidence interval.

#### Integration Performance

The Wasserstein critic demonstrated a strong advantage in local batch mixing. The critic achieved higher iLISI scores across all scenarios, indicating superior removal of local batch effects. Interestingly, global integration measures (ASW Batch) did not show the same effect; the discriminator performed comparably or slightly better (e.g., Lung and Pancreas). This discrepancy suggests that while the discriminator effectively aligns global distributions, the critic better resolves fine-grained batch structures within local cell neighborhoods.

#### Biological Conservation

A consistent trade-off was observed in biological conservation, with the aggressive batch mixing of the Wasserstein critic resulting in a reduction in cell-type purity. In terms of local neighborhood structure (cLISI), the critic exhibited higher scores compared to the discriminator, indicating that distinct cell types were less strictly separated in the local manifold. This pattern was corroborated by the global ASW Cell Type metric, where the critic yielded consistently lower scores, reflecting less compact clusters. The Adjusted Rand Index (ARI), which measures clustering agreement, showed similar patterns with the discriminator consistently outperforming the critic in the Pancreas and Immune datasets. The continuous topology metric (PAGA) showed a trend of cell type collapse in the critic compared to the discriminator, highlighting the over-corrective effects of the Wasserstein approach.

#### Sensitivity to Confounding Structure

We examined whether the correlation between batch and biological variables (Unbalanced scenarios) disproportionately affected either method. Contrary to the theoretical expectation that correlated confounding would universally degrade performance, we found no systematic pattern across datasets. For example, while the discriminator’s iLISI performance notably dropped in the Immune dataset, it actually improved in the Pancreas dataset. Similarly, the Wasserstein critic showed variable sensitivity: it maintained high performance in the Pancreas dataset but suffered a significant drop in the Immune and Lung datasets.

#### Summary

Overall, these results highlight a distinct behavior between the two adversaries. The discriminator acts as a conservative regularizer, preserving high within-cluster density (high ASW Cell Type) and cell type manifold (PAGA Spearman), but fails to fully integrate local neighborhoods (low iLISI). In contrast, the Wasserstein critic serves as a stronger adversary, forcing the encoder to thoroughly mix batches. This yields significant improvements in integration (iLISI) at the cost of a consistent reduction in cell-type compactness (ASW Cell Type) and topological location (PAGA Spearman), though the downstream effect on clustering (ARI) varies by biological context.

To visually confirm these quantitative trade-offs, we provide UMAP projections of the confounded gene expression data and deconfounded latent spaces in Supplementary Figures S3 and S4.

### Scalability and Stability in Multi-Batch Integration

To systematically evaluate the scalability of these methods, we incrementally increased the integration complexity from the binary baseline (|𝒱| = 2) up to the full dataset (|𝒱| = 9 for Pancreas, |𝒱| = 4 for Immune). We excluded the Lung dataset from this analysis as it contains only two batches. This experiment assesses whether the multi-headed Wasserstein formulation maintains its advantage as the number of alignment tasks grows (Figure 3).

**Figure 3.**
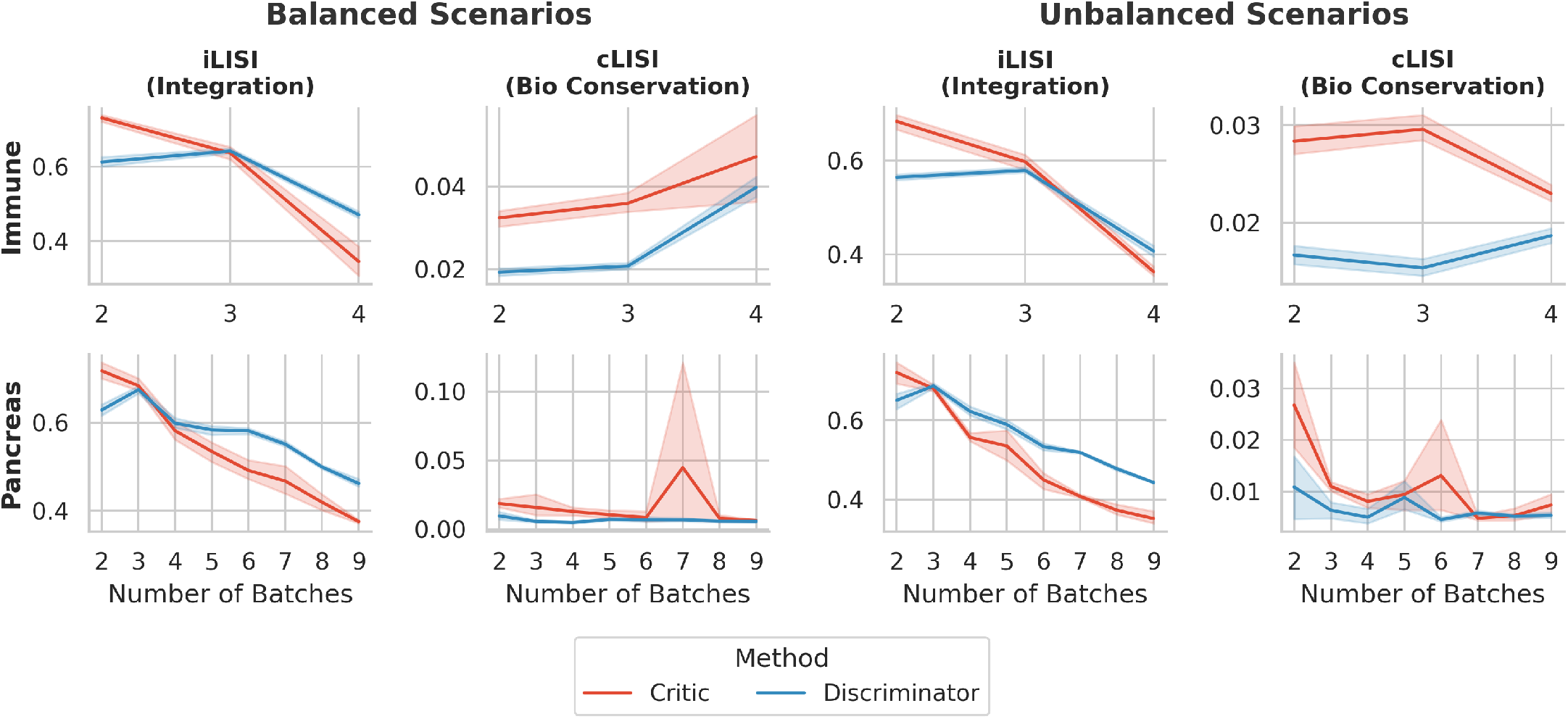
Impact of batch complexity on deconfounding performance. We evaluated the stability of the Wasserstein critic (Red line) and discriminator (Blue line) as the number of distinct batches increased (x-axis). The experiment was conducted across the Immune and Pancreas datasets under both Balanced and Unbalanced confounding scenarios. Performance is reported for **iLISI** (Integration, higher is better) and **cLISI** (Biological Conservation, lower is better). Solid lines represent the mean score, and error bars indicate the 95% confidence interval.

#### Performance Decay

As expected, integration difficulty increases with the number of batches. We observed a decrease in performance across all scenarios for both methods, reflecting the inherent challenge of aligning multiple disjoint distributions into a shared low-dimensional manifold.

#### The Discriminator Out-scales the Wasserstein Critic

The Wasserstein critic outperformed the discriminator on the iLISI metric in the binary case but scaled poorly as the number of batches increased. We hypothesize this highlights a critical trade-off in our reference-based multi-headed formulation. As |𝒱| increases, our critic must solve |𝒱| − 1 distinct transport problems to a single reference batch. In comparison, the discriminator optimizes a joint objective across the entire dataset, avoiding the topological bottleneck induced by forcing |𝒱|−1 independent distributions to map onto a single fixed target.

### Hyperparameter Sensitivity and Stability

A major practical limitation of adversarial deconfounding is the difficulty of tuning the regularization strength, *λ*. To evaluate the stability of the two objectives, we analyzed the impact of varying *λ* across orders of magnitude on both integration (iLISI) and conservation (cLISI) (Figure 4).

**Figure 4.**
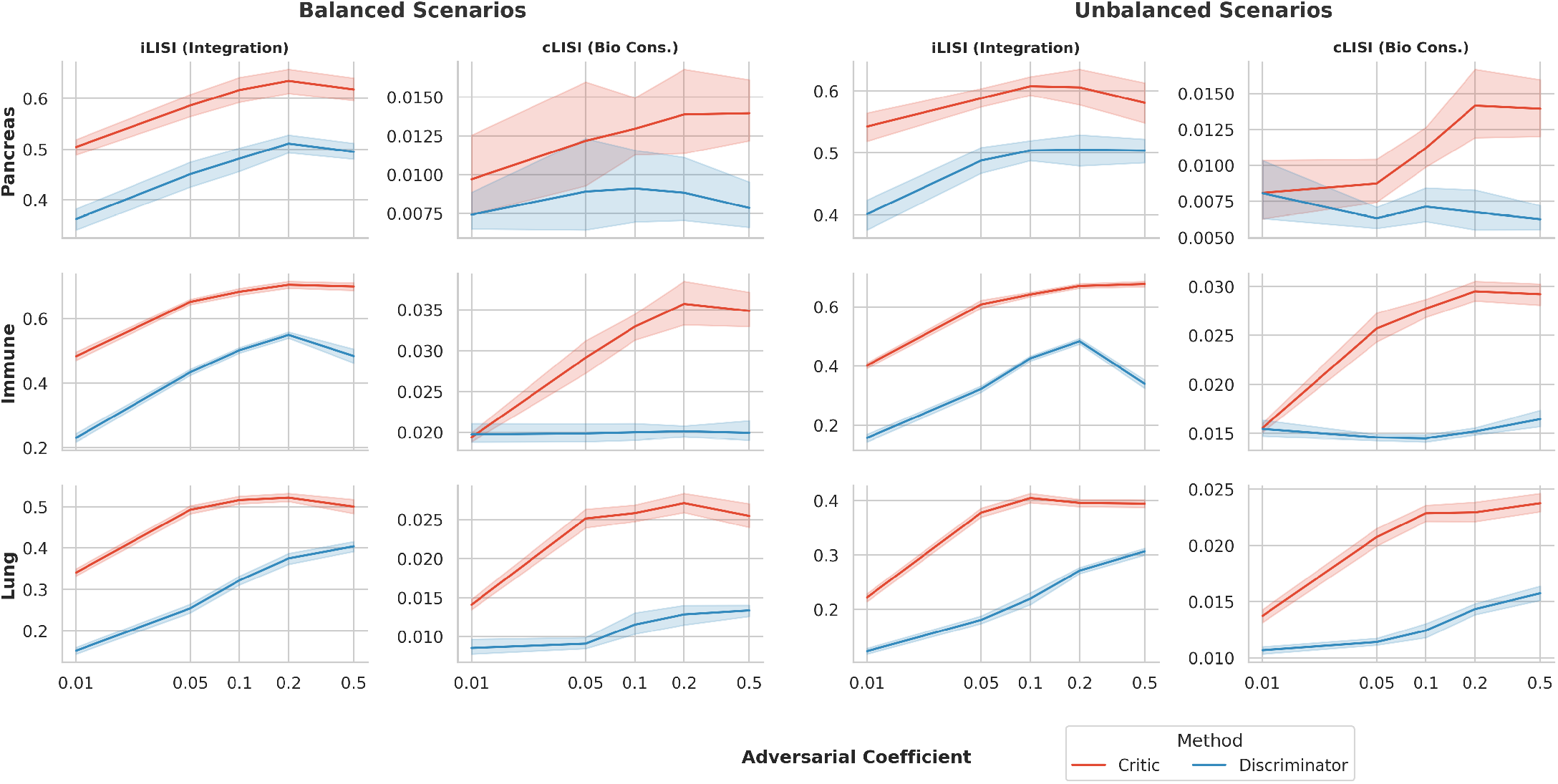
Sensitivity analysis of the adversarial coefficient across datasets. We evaluated the impact of varying the adversarial regularization strength (*λ*) on the performance of the Wasserstein critic (Red line) and discriminator (Blue line). The evaluation spans three datasets (Pancreas, Immune, Lung) under both Balanced and Unbalanced confounding scenarios. Performance is reported for **iLISI** (Integration, higher is better) and **cLISI** (Biological Conservation, lower is better). The x-axis represents the adversarial coefficient (*λ*) on a logarithmic scale. Solid lines represent the mean score across validation folds, and error bars indicate the 95% confidence interval.

#### Robustness to Regularization

Both the Wasserstein critic and the discriminator exhibit broad optimal operating windows on the iLISI metric, with the critic performing consistently higher than the discriminator. For the cLISI metric, the discriminator maintains a broad, flat curve that is highly insensitive to the choice of adversarial coefficient. In contrast, the Wasserstein critic’s cLISI score gradually increases as regularization strengthens, explicitly highlighting the continuous trade-off between aggressive batch integration and biological preservation.

### Impact of Reference Batch Selection

A unique architectural feature of our multi-headed critic is its alignment to a specific reference manifold. While this avoids the *O*(|𝒱|^2^) complexity of all-versus-all matching, it introduces a dependency on the quality of the chosen reference batch. To quantify this, we exhaustively trained models using every possible batch as the reference target and analyzed the variance in performance (Figure 5).

**Figure 5.**
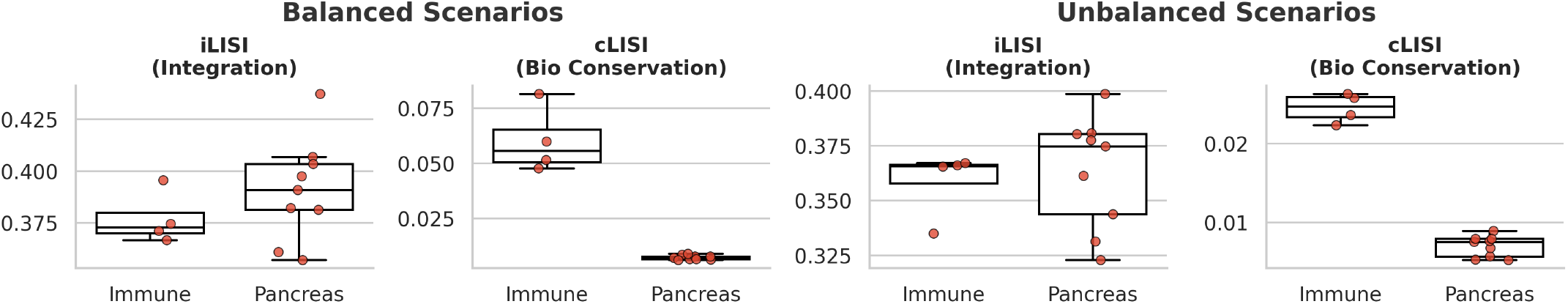
Sensitivity of the Wasserstein critic to reference batch selection. To assess the sensitivity to the reference manifold, we evaluated the performance of the Wasserstein critic using every possible batch as the reference target. The experiment spans the Pancreas and Immune datasets under both Balanced and Unbalanced confounding scenarios. Box plots summarize the distribution of performance scores, where each individual point (Red) represents the integrated result from a specific reference batch choice. Performance is reported for **iLISI** (Integration, higher is better) and **cLISI** (Biological Conservation, lower is better). A tighter distribution indicates that the method’s performance is robust regardless of which batch is chosen as the reference.

#### Sensitivity to Batch Completeness

The performance variance across reference batches reflects the underlying algorithmic sensitivity to reference density. As shown in Figure 5, significant outliers in iLISI performance emerge across both confounding scenarios. These outliers are strictly driven by the cell-type representation present in the target reference batch. Because the Wasserstein objective enforces a strict optimal transport plan, the model must map all cells from the source batch onto the reference manifold, even when the reference lacks corresponding cell populations. Consequently, when the chosen reference batch lacks representation of key cell types found in the query batches, the strict transport constraint forces the encoder to improperly project query-specific biological signals onto unrelated reference populations, effectively erasing true biological variance.

## Discussion

Adversarial deconfounding has become a cornerstone of deep learning-based batch correction, yet the choice of the adversarial objective itself has largely remained an implementation detail rather than a subject of rigorous study. In this work, we demonstrated that this choice is not merely technical but fundamental to the quality of integration. Our experiments reveal the following key results:

### The Wasserstein Critic is a Stronger Integrator than the JS-Discriminator

In scenarios with strong confounding or disjoint distributions, the discriminator frequently failed to mix local neighborhoods effectively. In contrast, the Wasserstein critic forced mixing even when batches were initially far apart.

### The Integration-Conservation Trade-off

While the Wasserstein critic significantly outperformed the discriminator in removing batch effects (higher iLISI), it incurred a consistent penalty in biological conservation (higher cLISI). Additionally, the continuous manifold (PAGA) was more warped in the case of the Wasserstein critic, forcing unrelated cell types together. Future work might explore adaptive weighting schemes, such as an annealing schedule that relaxes the adversarial penalty (*λ*_adv_) once a sufficiency threshold for mixing is reached. This could potentially capture the best of both worlds, achieving the aggressive mixing of the critic with the conservation of the discriminator.

### Structural Bottlenecks in Multi-Batch Scaling

While the Wasserstein critic excels in binary alignment, its integration performance degrades significantly faster than the discriminator as the number of experimental batches increases. Because this scalability drop occurs equally in both biologically balanced and unbalanced scenarios, it cannot be attributed to the reference batch lacking specific cell types. Instead, it reveals a fundamental architectural limitation of the reference-based multi-headed formulation. As the number of batches |𝒱| grows, the critic forces the network to solve |𝒱| − 1 independent, pairwise transport problems onto a single target geometry. This uncoordinated radial projection topology creates a geometric bottleneck, where multiple independent mappings inevitably distort the latent space to satisfy their individual transport constraints. In comparison, the discriminator optimizes a single, global |𝒱|-way classification task. By minimizing a joint divergence across the entire dataset simultaneously, the discriminator implicitly coordinates the latent geometry, smoothly aligning all batches toward a shared, centralized manifold without the compounding structural constraints of forcing multiple independent distributions to map onto a single anchor.

### From Reference Matching to Wasserstein Barycenters

The Wasserstein critic is further limited by the sparsity inherent to the reference matching induced by our multi-headed formulation, where a single reference batch acts as the central hub for |𝒱| − 1 target batches. While this topology avoids the quadratic complexity of all-vs-all matching, it is topologically fragile: if the reference is incomplete, the entire alignment suffers. The theoretical solution to this limitation is the *Wasserstein barycenter* (Agueh and Carlier, 2011). Unlike our current approach which fixes the center to an existing batch, a barycenter approach would learn a “virtual” center—the geometric median of all batch distributions. Implementing a barycenter objective would allow the model to align all batches to a shared, optimal manifold containing the union of all biological states, effectively combining the robust gradients of the Wasserstein critic with the global robustness of the discriminator.

### Limitations and Future Directions

To isolate the impact of the adversarial objective, we deliberately restricted our analysis to a single, state-of-the-art VAE backbone. While this controlled design was necessary to validate the mechanism, future work should evaluate whether these findings generalize to other architectures and models. Additionally, our current multi-headed formulation addresses a single categorical confounder. Extending this to handle multiple simultaneous confounders or continuous covariates would significantly broaden the applicability of the Wasserstein objective.

Our analysis indicates that the primary limitation of the Wasserstein objective in its isolated form is its propensity for biological over-correction. Future methodological developments must prioritize dynamic regularizers that prevent this topological collapse without sacrificing the robust optimization gradients provided by the Wasserstein metric.

## Supporting information

topology_matters_supplement

