## Supplementary material for "Topology Matters: The Trade-off Between Wasserstein Critics and Discriminators for Single-Cell Data Integration": topology_matters_supplement

#### Supplementary Note 1: Computational Efficiency

The Wasserstein critic requires multiple extra forward and backward passes during training. We investigated how this impacted total training time across our datasets.

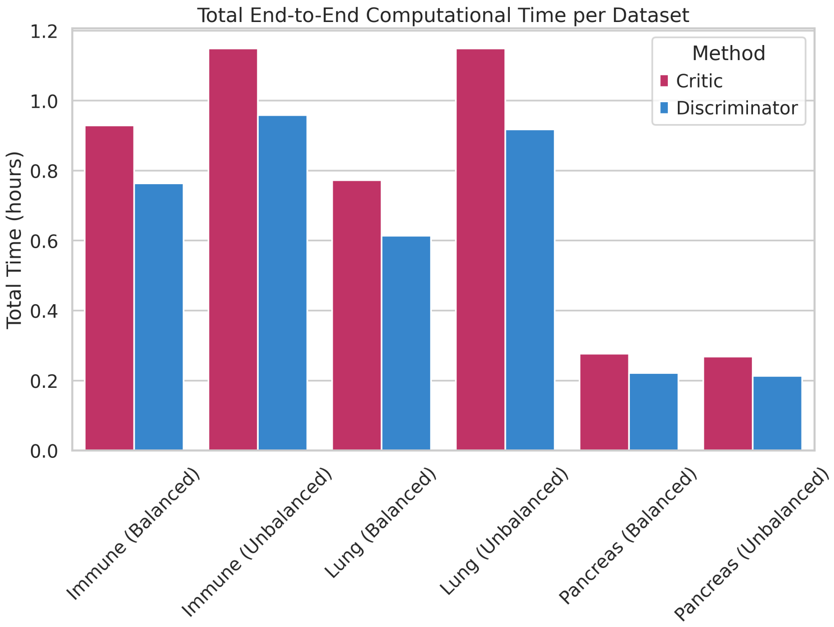


**Supplementary Figure 1: Computational Efficiency and Runtime.** Comparison of total training time across the Pancreas, Immune, and Lung datasets. The JS discriminator achieves faster convergence and requires significantly less computational overhead, as it does not require the computationally expensive gradient penalty computations inherent to the Wasserstein approach.

**Supplementary Table 1:** Detailed computational time differences per dataset and phase.

| Dataset Condition | Phase | Critic | Discriminator | Diff Seconds (Critic - Disc) | Diff Percentage |
|---|---|---|---|---|---|
| Immune (Balanced) | Final Outer Model | 113.6200 | 91.7980 | 21.8220 | 23.7718 |
| | Inner CV Fold | 37.0415 | 30.5373 | 6.5041 | 21.2990 |
| Immune (Unbalanced) | Final Outer Model | 138.3800 | 114.4960 | 23.8840 | 20.8601 |
| | Inner CV Fold | 45.9227 | 38.3869 | 7.5357 | 19.6310 |
| Lung (Balanced) | Final Outer Model | 92.9860 | 72.9340 | 20.0520 | 27.4934 |
| | Inner CV Fold | 30.8929 | 24.5988 | 6.2941 | 25.5872 |
| Lung (Unbalanced) | Final Outer Model | 137.2920 | 108.5220 | 28.7700 | 26.5108 |
| | Inner CV Fold | 46.0029 | 36.8245 | 9.1784 | 24.9247 |
| Pancreas (Balanced) | Final Outer Model | 31.4280 | 25.4040 | 6.0240 | 23.7128 |
| | Inner CV Fold | 11.1949 | 8.9272 | 2.2677 | 25.4025 |
| Pancreas (Unbalanced) | Final Outer Model | 30.5960 | 24.2800 | 6.3160 | 26.0132 |
| | Inner CV Fold | 10.8571 | 8.6091 | 2.2480 | 26.1120 |

#### Supplementary Note 2: Dataset Composition

To evaluate the limits of adversarial integration, we utilized three highly complex single-cell transcriptomic datasets (Pancreas, Lung, Immune) characterized by severe batch effects, varying sequencing depths, and extreme class imbalances. Supplementary Table 2 details the hierarchical composition of all three datasets.

**Supplementary Table 2:** Hierarchical composition of datasets by Batch and Cell Type.

| **Pancreas Dataset** | | **Lung Dataset** | | **Immune Dataset** | |
|---|---|---|---|---|---|
| **Batch / Cell Type** | **N** | **Batch / Cell Type** | **N** | **Batch / Cell Type** | **N** |
| **CEL-Seq** | **1,004** | **10x Chromium** | **22,771** | **10X** | **8,829** |
| acinar | 228 | B cell | 103 | CD4+ T cells | 4,312 |
| activated_stellate | 19 | Basal 1 | 1,972 | CD8+ T cells | 578 |
| alpha | 191 | Basal 2 | 3,072 | CD14+ Monocytes | 1,501 |
| beta | 161 | Ciliated | 2,722 | CD16+ Monocytes | 271 |
| delta | 50 | Dendritic cell | 1,367 | CD20+ B cells | 409 |
| ductal | 327 | Endothelium | 628 | Megakaryocyte progenitors | 14 |
| endothelial | 5 | Fibroblast | 529 | Monocyte-derived dendritic cells | 82 |
| epsilon | 1 | Ionocytes | 46 | NK cells | 973 |
| gamma | 18 | Lymphatic | 146 | NKT cells | 649 |
| macrophage | 1 | Macrophage | 4,614 | Plasmacytoid dendritic cells | 40 |
| mast | 1 | Mast cell | 229 | **smart-seq2** | **1,022** |
| quiescent_stellate | 1 | Neutrophil_CD14_high | 1,626 | CD14+ Monocytes | 145 |
| schwann | 1 | Neutrophils_IL1R2 | 472 | CD16+ Monocytes | 172 |

**Supplementary Table 2:** Hierarchical composition of datasets by Batch and Cell Type (Continued)

| **Pancreas Dataset** | | **Lung Dataset** | | **Immune Dataset** | |
|---|---|---|---|---|---|
| **Batch / Cell Type** | **N** | **Batch / Cell Type** | **N** | **Batch / Cell Type** | **N** |
| **CEL-Seq2** | **2,285** | Secretory | 1,753 | Monocyte-derived dendritic cells | 534 |
| acinar | 274 | T/NK cell | 616 | Plasmacytoid dendritic cells | 171 |
| activated_stellate | 90 | Type 2 | 2,876 | **10x Chromium v2** | **12,928** |
| alpha | 843 | **Drop-Seq** | **6,485** | CD4+ T cells | 3,762 |
| beta | 445 | B cell | 916 | CD8+ T cells | 1,255 |
| delta | 203 | Ciliated | 313 | CD10+ B cells | 207 |
| ductal | 258 | Endothelium | 219 | CD14+ Monocytes | 1,449 |
| endothelial | 21 | Fibroblast | 147 | CD16+ Monocytes | 190 |
| epsilon | 4 | Lymphatic | 128 | CD20+ B cells | 918 |
| gamma | 110 | Macrophage | 2,036 | Erythrocytes | 1,502 |
| macrophage | 15 | Mast cell | 488 | Erythroid progenitors | 463 |
| mast | 6 | Secretory | 295 | HSPCs | 445 |
| quiescent_stellate | 12 | T/NK cell | 749 | Megakaryocyte progenitors | 235 |
| schwann | 4 | Type 1 | 297 | Monocyte progenitors | 428 |
| **Fluidigm C1** | **638** | Type 2 | 897 | Monocyte-derived dendritic cells | 214 |
| acinar | 21 | | | NK cells | 565 |
| activated_stellate | 16 | | | NKT cells | 1,040 |
| alpha | 239 | | | Plasma cells | 111 |
| beta | 258 | | | Plasmacytoid dendritic cells | 144 |
| delta | 25 | | | **10x Chromium v3** | **10,727** |
| ductal | 36 | | | CD4+ T cells | 2,937 |
| endothelial | 14 | | | CD8+ T cells | 350 |
| epsilon | 1 | | | CD14+ Monocytes | 3,388 |
| gamma | 18 | | | CD16+ Monocytes | 364 |
| macrophage | 1 | | | CD20+ B cells | 1,546 |
| mast | 3 | | | HSPCs | 28 |
| quiescent_stellate | 1 | | | Megakaryocyte progenitors | 21 |

**Supplementary Table 2:** Hierarchical composition of datasets by Batch and Cell Type (Continued)

| **Pancreas Dataset** | | **Lung Dataset** | | **Immune Dataset** | |
|---|---|---|---|---|---|
| **Batch / Cell Type** | **N** | **Batch / Cell Type** | **N** | **Batch / Cell Type** | **N** |
| schwann | 5 | | | | 182 |
| | | | | Monocyte-derived dendritic cells | |
| **inDrop-Seq V1** | **1,937** | | | NK cells | 756 |
| acinar | 110 | | | NKT cells | 1,056 |
| activated_stellate | 51 | | | Plasma cells | 18 |
| alpha | 236 | | | Plasmacytoid dendritic cells | 81 |
| beta | 872 | | | | |
| delta | 214 | | | | |
| ductal | 120 | | | | |
| endothelial | 130 | | | | |
| epsilon | 13 | | | | |
| gamma | 70 | | | | |
| macrophage | 14 | | | | |
| mast | 8 | | | | |
| quiescent_stellate | 92 | | | | |
| schwann | 5 | | | | |
| t_cell | 2 | | | | |
| **inDrop-Seq V2** | **1,724** | | | | |
| acinar | 3 | | | | |
| activated_stellate | 81 | | | | |
| alpha | 676 | | | | |
| beta | 371 | | | | |
| delta | 125 | | | | |
| ductal | 301 | | | | |
| endothelial | 23 | | | | |
| epsilon | 2 | | | | |
| gamma | 86 | | | | |
| macrophage | 17 | | | | |
| mast | 9 | | | | |
| quiescent_stellate | 22 | | | | |
| schwann | 6 | | | | |
| t_cell | 2 | | | | |
| **inDrop-Seq V3** | **3,605** | | | | |
| acinar | 843 | | | | |
| activated_stellate | 100 | | | | |
| alpha | 1,130 | | | | |
| beta | 787 | | | | |
| delta | 161 | | | | |
| ductal | 376 | | | | |
| endothelial | 92 | | | | |
| epsilon | 2 | | | | |
| gamma | 36 | | | | |

**Supplementary Table 2:** Hierarchical composition of datasets by Batch and Cell Type (Continued)

| **Pancreas Dataset** | | **Lung Dataset** | | **Immune Dataset** | |
|---|---|---|---|---|---|
| **Batch / Cell Type** | **N** | **Batch / Cell Type** | **N** | **Batch / Cell Type** | **N** |
| macrophage | 14 | | | | |
| mast | 7 | | | | |
| quiescent_stellate | 54 | | | | |
| schwann | 1 | | | | |
| t_cell | 2 | | | | |
| **inDrop-Seq V4** | **1,303** | | | | |
| acinar | 2 | | | | |
| activated_stellate | 52 | | | | |
| alpha | 284 | | | | |
| beta | 495 | | | | |
| delta | 101 | | | | |
| ductal | 280 | | | | |
| endothelial | 7 | | | | |
| epsilon | 1 | | | | |
| gamma | 63 | | | | |
| macrophage | 10 | | | | |
| mast | 1 | | | | |
| quiescent_stellate | 5 | | | | |
| schwann | 1 | | | | |
| t_cell | 1 | | | | |
| **SMARTer** | **1,492** | | | | |
| alpha | 886 | | | | |
| beta | 472 | | | | |
| delta | 49 | | | | |
| gamma | 85 | | | | |
| **Smart-Seq V2** | **2,394** | | | | |
| acinar | 188 | | | | |
| activated_stellate | 55 | | | | |
| alpha | 1,008 | | | | |
| beta | 308 | | | | |
| delta | 127 | | | | |
| ductal | 444 | | | | |
| endothelial | 21 | | | | |
| epsilon | 8 | | | | |
| gamma | 213 | | | | |
| macrophage | 7 | | | | |
| mast | 7 | | | | |
| quiescent_stellate | 6 | | | | |
| schwann | 2 | | | | |

### Supplementary Note 3: Visualization of Confounded Gene Expression Data

To gain an intuitive understanding of the different latent spaces, we compared the UMAP results from the Pancreas dataset before and after deconfounding with both the discriminator and critic.

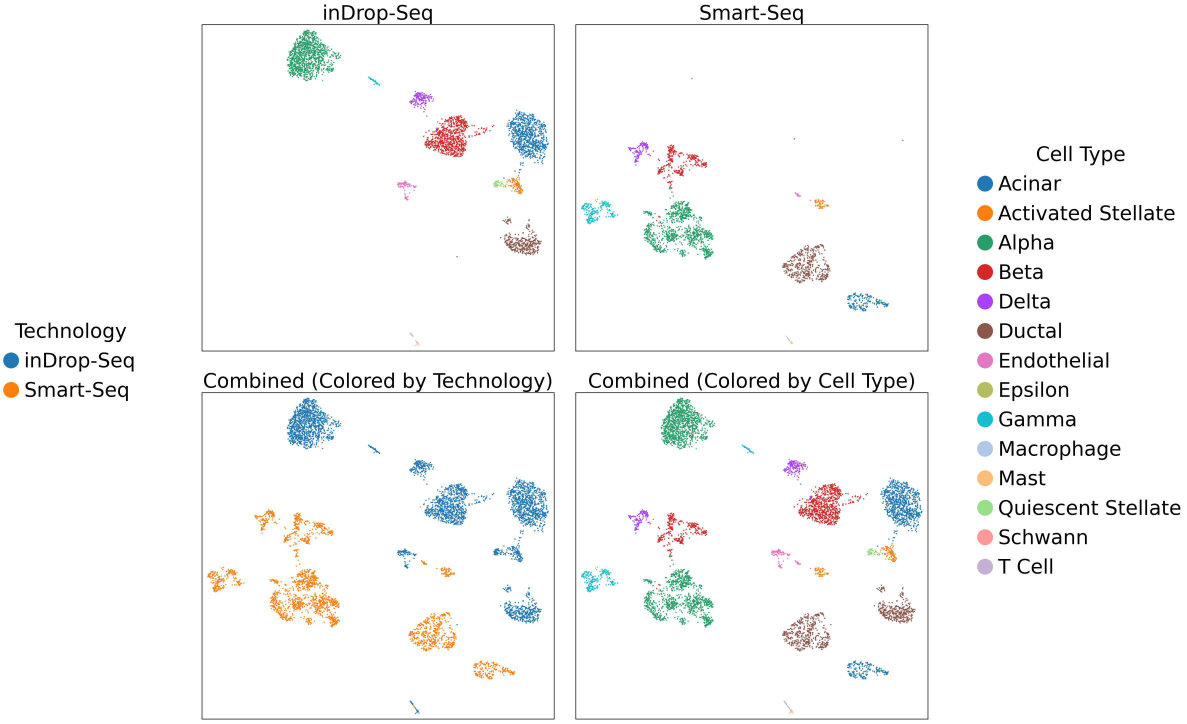


**Supplementary Figure 2: Pancreas Dataset before Deconfounding.** Visualization of the Pancreas dataset. **Top row:** UMAP of inDrop-Seq and Smart-Seq datasets colored by cell type. **Bottom left:** Combined UMAP of inDrop-Seq (blue) and Smart-Seq (yellow) colored by technology. The two technologies are strongly separated. **Bottom right:** Combined UMAP of inDrop-Seq and Smart-Seq colored by cell type. There are two islands for each cell type corresponding to the two different technologies.

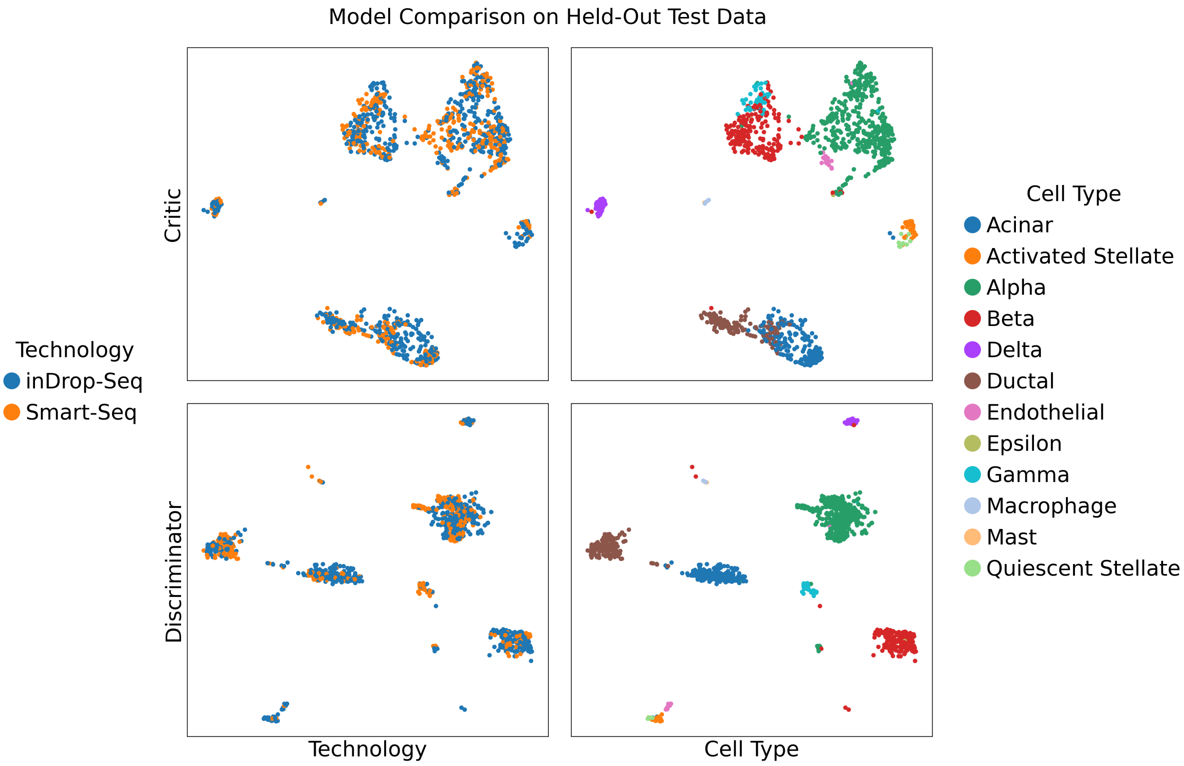


**Supplementary Figure 3: Pancreas Dataset after Deconfounding.** Visualization of the Pancreas dataset latent space for both the critic and discriminator. **Top left:** The critic latent space colored by technology. The two technologies are strongly overlapped. **Top right:** The critic latent space colored by cell type. The cell populations blend together along the boundaries of the respective clusters. **Bottom left:** The discriminator latent space colored by technology. The two technologies are integrated in most areas; however, there are some distinct regions that remain unmixed. **Bottom right:** The discriminator latent space colored by cell type. The different cell type clusters are very distinct.
